# Insights into Parkinson’s disease from computational models of the basal ganglia

**DOI:** 10.1101/260992

**Authors:** Mark D. Humphries, Jose Obeso, Jakob Kisbye Dreyer

## Abstract

Movement disorders arise from the complex interplay of multiple changes to neural circuits. Successful treatments for these disorders could interact with these complex changes in myriad ways, and as a consequence their mechanisms of action and their amelioration of symptoms are incompletely understood. Using Parkinson’s disease as a case-study, we review here how computational models are a crucial tool for taming this complexity, across causative mechanisms, consequent neural dynamics, and treatments. For mechanisms, we review models that capture the effects of losing dopamine on basal ganglia function; for dynamics, we discuss models that have transformed our understanding of how beta-band (15-30 Hz) oscillations arise in the parkinsonian basal ganglia. For treatments, we touch on the breadth of computational modelling work trying to understand the therapeutic actions of deep brain stimulation. Collectively, models from across all levels of description are providing a compelling account of the causes, symptoms, and treatments for Parkinson’s disease.

## Introduction

The basal ganglia are implicated in a wide range of movement disorders, especially Parkinson’s disease, Huntington’s disease, and dystonia. The causes and progression of these disorders are complex, arising from the interplay between multiple changes to neural circuits, and between those changes and consequent compensatory mechanisms in the brain. The link between these changes and the resultant overt clinical manifestations is incompletely understood. Available treatments are of limited efficacy and often have poorly defined mechanisms of action. As we aim to show here, one route out of this thicket is to use computational models as a guide to our often faulty intuition

Parkinson’s disease exemplifies these issues, and will be our focus here. Classically, the onset of its cardinal signs (bradykinesia/akinesia, tremor, and rigidity) correlates with the loss of midbrain dopamine neurons projecting to the dorsal striatum, and especially the putamen in humans. This has long suggested a role for the basal ganglia in motor control, and pointed to aberrant basal ganglia dynamics as the root cause of the cardinal motor features. But the basal ganglia are a densely connected web of nuclei (Figure 1A). Any change to one neural population in the basal ganglia will have effects that ripple throughout its components; changes to more than one population are impossible to predict with any confidence. Consequently, there is a need for computational models of the parkinsonian basal ganglia to aid our understanding.

**Figure 1:**
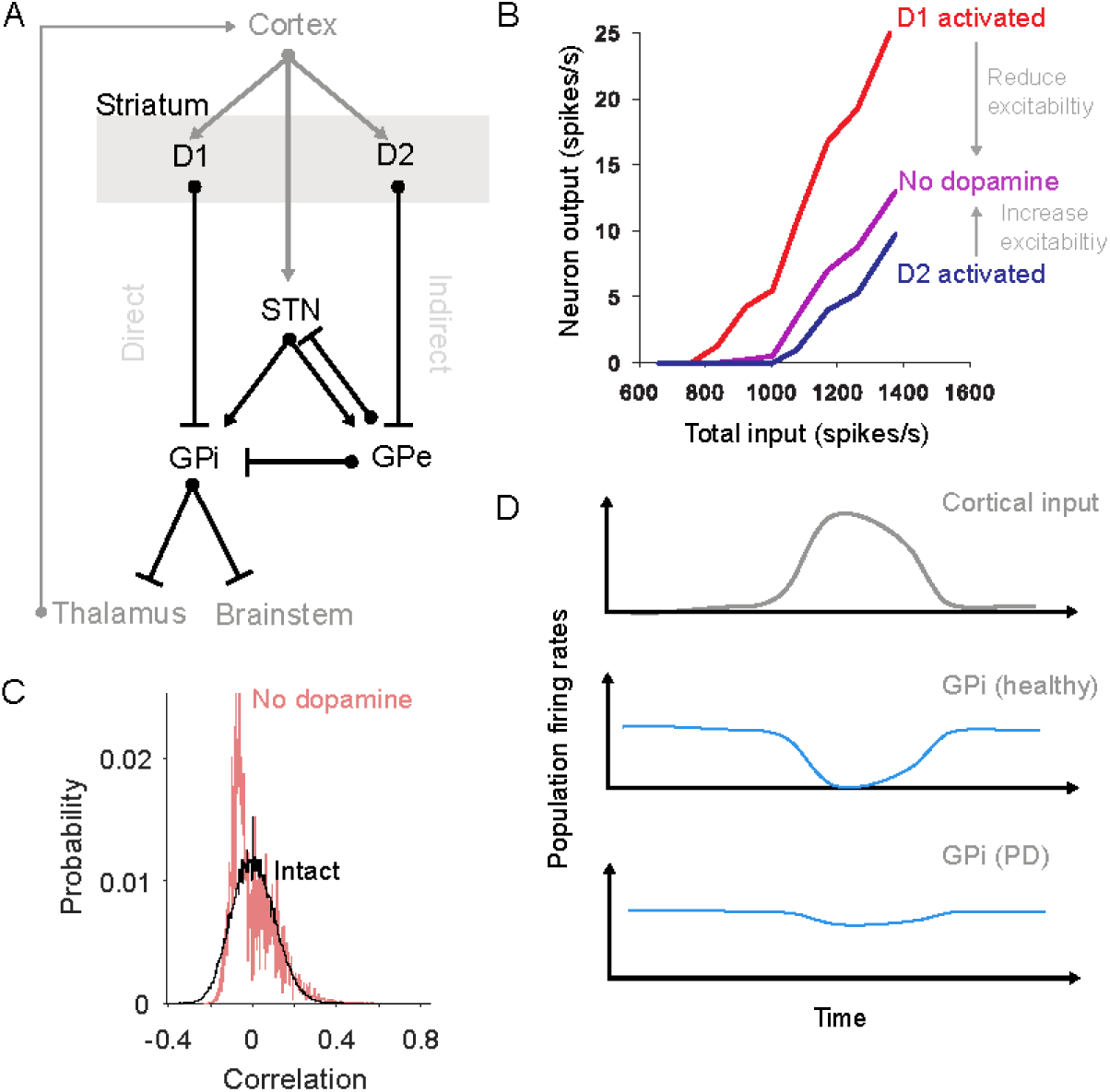
Consequences of dopamine depletion in the striatum. **A** The basal ganglia nuclei and main connections (bar: inhibitory; arrow: excitatory). Striatum divides into two projection neurons populations expressing D1 and D2 type receptors for dopamine. Routes from the D1 and D2 populations to the output nuclei (GPi) have historically been labelled the “direct” and “indirect” pathways. For clarity, we omit here some pathways, including the projection from GPe to the striatum, and the local connections within the striatum. STN: subthalamic nucleus; GPe: globus pallidus, external segment; GPi: globus pallidus, internal segment. **B** Single neuron models predict that dopamine differentially affects the excitability of D1 and D2-receptor expressing projection neurons. Consequently, dopamine depletion reduces excitability of D1-expressing neurons, and increases the excitability of D2-expressing neurons. Redrawn from [13]. **C** Distributions of the correlations between spontaneous neuron firing in intact and dopamine-depleted striatal network models [16]. The models predict that the spontaneous activity of the intact network is sparse, irregular and uncorrelated; but that dopamine depletion creates spontaneous activity that is anti-correlated (negative correlations). Such changes within the striatum’s dynamics are in addition to any changes to the drive or synchrony of its cortical input caused by dopamine depletion. **D** Schematics showing how network models of the basal ganglia predict a breakdown of action selection in Parkinson’s disease [cartoon of the results in e.g. 18, 20, 23]. Under normal conditions, a phasic input to the basal ganglia from cortex (top) produces (middle) a transient suppression of activity in a small group of GPi neurons (blue); this transient suppression of inhibition allows “selection” to occur by disinhibiting the thalamic and brainstem targets of these GPi neurons. Following dopamine depletion (bottom), the GPi activity barely changes in response to the same input, thus slowing or preventing action selection.

There is a rich history of computational modelling of the basal ganglia’s dynamics in their healthy state, and of their changes under parkinsonian conditions [1–3]. Here we review key insights from such modelling in understanding neural mechanisms of the disease and its symptoms, how its aberrant neural dynamics arise, and how its treatments work.

### Consequences of dopamine depletion in the striatum

We begin with the striatum. The striatum contains the densest expression of dopamine receptors in the vertebrate brain [4, 5]. The putamen is also the target of the most vulnerable set of midbrain dopamine neurons in Parkinson’s disease [6]. Thus anyone interested in understanding the causes of motor deficits in Parkinson’s disease is naturally drawn to understanding the effects of dopamine depletion on the striatum.

The effects of dopamine receptor activation on a single striatal neuron are subtle and complex [7–9]. Experimental studies of the striatal projection neuron have established how dopamine receptor activation down- and up-regulates a suite of ion channels, both inside and outside of synapses [7], to control the excitability of the neuron. Understanding the interplay between all these effects has required detailed models of single projection neurons that synthesise the findings of many experimental studies [10–12]. One contribution of these models has been to provide an intuitive picture of the net effect of activating either D1 or D2 type dopamine receptors, and show that these effects cause D1 and D2 expressing projection neurons to respond differently to excitatory input [13]. They thus predict that the net effect of dopamine depletion is to change the excitability of projection neurons in opposite directions depending on their expression of D1 or D2 type dopamine receptors (Figure 1B). Such models have shown that intuitive arguments about Parkinson’s disease changing the balance of the direct and indirect pathways [14, 15] have a biophysical basis.

Consistent with the changes to individual neuron dynamics, models of the whole striatal network predict that dopamine depletion profoundly disrupts its normal dynamics. One study suggests that dopamine-depletion increases the spontaneous correlations between projection neuron firing [16]; such spontaneous correlated activity could potently block cortical input from being transmitted to striatum and onto the rest of the basal ganglia (Figure 1C). Another study suggests that dopamine depletion alters the balance of D1 and D2 projection neuron activity by changing the output of the sparse but powerful inhibitory interneurons [17]. Both point to the need to understand how network-scale changes in Parkinson’s disease arise from the cumulative effects of dopamine depletion on individual neurons.

At a broader scale still, models of the whole basal ganglia network have sought to understand the knock-on effects of the depletion of dopamine in the striatum. Many such models test the theory that the basal ganglia’s normal function is to perform action selection through the interaction of the direct and indirect pathways [18, 19]. Dopamine depletion in these models disrupts the balance of the two striatal output pathways, leading to action selection deficits [20-22]. The imbalance could arise either through direct effects on neural excitability, as reviewed above, or through aberrant cortico-striatal plasticity that follows dopamine depletion [21]. Other models suggest a disruption between the balance of direct and cortico-subthalamic (or “hyperdirect”) pathways by dopamine depletion [21,23]. Whatever the mechanism, all these models predict a jamming of basal ganglia output, because the direct pathway can no longer induce sufficient inhibition of the basal ganglia output (Figure 1D). This lost ability to voluntarily select actions is consistent with the akinetic and bradykinetic features of Parkinson’s disease.

### Modelling the effect of losing dopamine neurons on the concentration of dopamine and on D1 and D2 signalling

While the above models seek the consequences of dopamine depletion on the dynamics of the striatum and wider basal ganglia network, others have pursued the equally profound question of how the loss of midbrain dopamine cells creates a complex landscape of adaptations to that loss and subsequent changes in dopamine dynamics during the development of Parkinson’s disease. They suggest that a more nuanced approach is needed to understand the effects of dopamine loss on striatum, and thus on basal ganglia function.

One of the simplest but potentially far-reaching predictions of these release models is the phenomenon of passive stabilisation [24]. Kinetic models of dopamine release are often used to simulate the interplay between dopamine release and re-uptake [25, 26]. The loss of dopamine terminals in Parkinson’s disease simultaneously causes a reduction in vesicular release of dopamine and in its re-uptake. Kinetic models predict that the loss of release and of re-uptake are exactly balanced, and as a consequence there is little change in steady-state dopamine tone following the loss of dopamine terminals [24, 27]. This passive stabilisation of dopamine tone is even more robust if the effects of dopamine autoreceptors on the terminals are simulated, because the autoreceptors act as negative feedback on vesicle release [28]. Passive stabilisation of dopamine tone is also seen in detailed reaction-diffusion models of dopamine release that emulate the volume transmission of dopamine throughout a three-dimensional region of the striatum (Figure 2A) [29–31]. Thus several lines of modelling predict that passive stabilisation maintains a normal dopamine tone despite the loss of dopamine neurons (Figure 2B-C).

**Figure 2:**
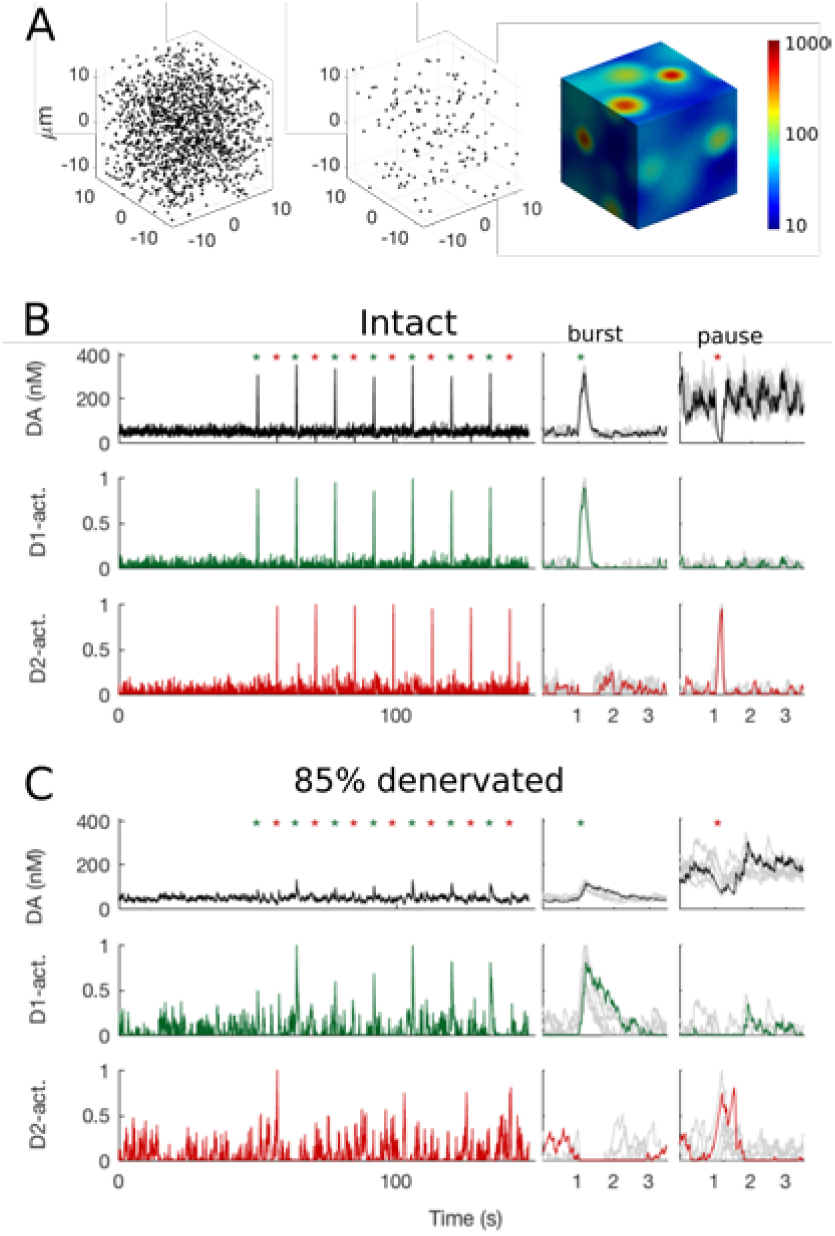
Modelling of dopamine volume transmission after cell loss. **A** Modelling dopamine release, diffusion, and re-uptake in volumes of the striatum [29–31]. For a 20 micrometre cube of simulated striatum, we show the simulated density of dopamine release sites in healthy tissue (left) and after losing 90% of dopamine terminals (middle). Simulating dopamine release from all sites results in heterogeneous dopamine concentration throughout the volume (color-scale in nM). **B** Simulated effects of bursts and pauses in dopamine cell firing on dopamine concentration (top). The average concentration of dopamine (dopamine tone) is maintained around 40 nM throughout. Alternating bursts (green asterisks) and pauses (red asterisk) in spike firing cause phasic peaks and dips in dopamine concentration. These consequently activate dopamine receptor-dependent intracellular signals via D1 (green, middle) and D2 (red, bottom) receptors: D1 receptors read-out peaks; D2 receptors read-out dips. Insets show averaged responses around each burst or pause (individual responses in grey). **C** As panel B, for a volume with 85% loss of dopamine release sites. Despite the severe loss, passive stabilisation means that the average concentration of dopamine is unchanged. By contrast, changes in dopamine concentration during bursts and pauses are now barely discernible from random fluctuations in the background dopamine tone (top). Consequently, the model predicts that D1 and D2 receptor dependent signals will be similar between periods of background tone and periods of phasic events (middle and bottom).

Passive stabilisation seemingly provides a simple hypothesis [24, 27] for why the cardinal signs of Parkinson’s disease develop after substantial loss of midbrain dopamine neurons. The motor features of Parkinson’s disease are often thought to be a consequence of a gradual loss of dopamine tone; from this perspective, the delayed appearance of the motor signs of Parkinson’s disease could be explained as a consequence of this passive stabilisation of dopamine tone.

This cannot though be the whole story. Whereas post-mortem and imaging studies estimate that a loss of dopaminergic neurons and terminals on the order of 50% correlates with the emergence of parkinsonian motor features [6, 32], detailed volume transmission models predict that dopamine tone would still be stabilised with a nearly complete loss of dopamine neurons and terminals [31, 33]. As an example, Figure 2C shows these models predict that losing 85% of the dopamine terminals has no detectable effect on dopamine tone. Volume transmission models instead predict that the onset of the cardinal signs of Parkinson’s disease is a consequence of the degraded *phasic* variations in dopamine concentration against this stabilised background tone.

Phasic variations in dopamine concentration arise because dopamine neurons deviate around their constant spontaneous activity with brief bursts or pauses (Figure 2B). These phasic changes in firing can be elicited by unexpected rewards and events [34–36], which may act as a teaching signal that controls synaptic plasticity in the striatum [37–40]. Phasic changes in both the firing of dopamine neurons in the lateral substantia nigra pars compacta [S41] and the activity of dopaminergic axons in the dorsal striatum [S42] also correlate with movement. Volume transmission models of dopamine in the intact striatum show that these bursts and pauses in activity are translated with high fidelity into phasic increases and decreases in dopamine concentration [29, 30] (Figure 2B). However, they also show that the loss of dopamine terminals inevitably decreases how effectively bursts and pauses of activity can change dopamine concentration (Figure 2C) [31].

These models predict that because the dopamine tone is passively stabilised, but phasic changes in dopamine are blunted, so the phasic variations become more difficult to distinguish from mere fluctuations in dopamine tone as more dopamine neurons are lost. This reduction in the signal-to-noise ratio of phasic dopamine becomes notable at moderate loss of dopamine terminals, consistent with the predicted scale of loss of dopamine neurons and terminals at which parkinsonian motor features are first observed [6, 32, S43]. The models predict that it finally becomes impossible to distinguish phasic events from random fluctuations in baseline dopamine tone at around 80% loss of terminals [31] (Figure 2C). In addition, as the magnitude of the phasic variations in dopamine decrease with the loss of dopamine neurons, so the dopamine receptors are likely to increase their sensitivity to compensate. The combined effect of reduced signal-to-noise ratio and increased receptor sensitivity is thus predicted to be aberrant activation of the D1 and D2 receptors, uncoupled from external events.

This aberrant activation has potentially many functional consequences. On the one hand, dopamine-mediated teaching signals would be missed, or random fluctuations treated as teaching signals, both leading to aberrant synaptic plasticity in striatum. On the other, if phasic variations in dopamine act as a signal to initiate movement [S41,S42], then losing the ability to detect phasic changes would slow or impair the initiation of movement.

The volume transmission models also predict that the breakdown of passive stabilisation after massive loss of terminals is not homogeneous, as it creates isolated volumes of tissue that contain no dopamine innervation [31]. These dopamine voids are predicted to create local imbalances in striatum, as neurons within neighbouring regions of intact and depleted dopamine interact. The volume transmission models thus point to a dual faced nature of dopamine denervation in Parkinson’s disease, first through loss of phasic dopamine dynamics and later through the creation of dopamine voids, raising the possibility that symptoms and signs arise by these different mechanisms acting concurrently in different areas of the striatum.

A key insight from computational models is that striatal neurons expressing the D1 and D2 receptors are differentially sensitive to the loss of dopamine terminals [11]. Modelling suggests that D1 receptors will respond strongly to phasic peaks in dopamine concentration, driven by bursts of spikes at up to 30 Hz; by contrast, models predict that D2 receptors respond to phasic dips in dopamine concentration, driven by pauses in the 4Hz spontaneous rate of spiking [11, 12]. As these downward deflections in firing rate are bound below by zero, so the D2 receptors have a worse signal-to-noise ratio for detecting phasic dopamine events even in the intact striatum (Figure 2B). Consequently, models predict that D2 receptors will be more sensitive to the loss of dopamine terminals [31, 33,S41]. These predictions are consistent with the early remodelling of D2-expressing striatal projection neurons in rodent models of Parkinson’s disease [S44].

Detailed kinetic models also allow us a better understanding of L-DOPA’s therapeutic actions and side-effects. Network models of the striatum simulate L-DOPA as a global increase in dopamine [19, 21, S45]. Such models predict that a side-effect of L-DOPA would be to raise dopamine tone above normal levels in striatal regions that had not lost dopamine terminals. Detailed release models more accurately simulate L-DOPA as an increase in the number of dopamine molecules released by vesicles in remaining terminals [31, 33]. In direct contrast to the network models, these release models predict little effect in healthy regions of the striatum due to autoreceptors regulating release at dopamine terminals. In dennervated regions, release models predict that L-DOPA’s increase of vesicles allows both partial restoration of dopamine tone and improved separation of phasic dopamine release from background tone [31]. As these models also capture homeostatic changes in receptor sensitivity, a future avenue for work could be to examine the consequences of on-medication periods for dopamine signalling in subsequent off-medication periods, in order to dig deeper into the origins of L-DOPA’s side-effects.

### Mechanisms of Parkinsonian neural oscillations

Our clearest glimpse of the neural dynamics in the parkinsonian basal ganglia has come from recordings obtained during surgery to implant electrodes for deep brain stimulation. Recordings from such electrodes placed in the subthalamic nucleus have revealed a prominent 15-30 Hz “beta-band” oscillation in the local field potential, a signature of co-ordinated synaptic activity [S46, S47]. The strength of beta-band oscillation correlates with motor-deficit severity, is suppressed by dopamine-replacement medication, and the magnitude of its suppression is correlated with the degree of improvement in movement [S48,S49]. Consequently, the questions of how and where such an oscillation arises have driven a rich vein of modelling work.

Modellers have pursued three broad hypotheses for the origin of the beta-band oscillation. The first and most popular has been the negative feedback loop between the excitatory subthalamic nucleus (STN) and the inhibitory external globus pallidus (GPe). Modellers have long found this loop intriguing because delayed negative feedback loops are a classic electrical circuit design for oscillators; perhaps unsurprisingly, models of the STN-GPe loop have been shown to generate oscillations under a wide range of conditions, both healthy and parkinsonian [20, S50–S52].

By what mechanisms does this loop produce oscillations under parkinsonian conditions of dopamine depletion? Models have uncovered multiple mechanisms that can cause this loop to shift from stable to oscillatory activity [22], but two have been prominently explored because they could plausibly follow from the loss of dopamine. One mechanism is the strengthening of the effect of input from D2-receptor striatal projection neurons to the pallidum [S50], possibly due to the increased excitability of D2 projection neurons (which is a predicted consequence of dopamine loss, as noted above). The second mechanism is the strengthening of the connections between STN and GPe [20], possibly because pre-synaptic D2 receptors that prevent transmitter release in these nuclei are no longer activated after dopamine depletion [20]. Either alone or in combination, both these mechanisms switch the STN-GPe loop from stable to oscillating.

While there are many routes to making the STN-GPe loop oscillate, models have shown that specifically producing oscillations in the beta-band requires a more limited set of conditions. Analytical models from Bogacz and colleagues [S53,S54] showed that beta-band oscillations can arise if the total delays in transmission between the STN and GPe are set within a narrow range. Biophysical models of the STN-GPe loop also require setting specific transmission delays to obtain beta-band oscillations [S55]. This raised the question of whether the transmission delays in the real primate basal ganglia meet these conditions. Recently, building on their detailed model of primate basal ganglia [S56], Lienard and colleagues [S57] searched for the set of transmission delays that allowed their model to replicate a range of electrophysiological data in healthy primates. With their found set of delays in hand they then made their model parkinsonian, by increasing the strength of connections between STN and GPe, and beta-band oscillations emerged. Models thus show that beta-band oscillations in the basal ganglia of primates can emerge from the STN-GPe loop. But a challenge to this idea is that while rodent models of Parkinson’s also show beta-band oscillations within STN [S58], computational models suggest the STN-GPe loop in rodents naturally oscillates at higher frequencies [20]; this suggests beta-band oscillations in rodents have their origin outside the STN-GPe loop.

Modellers have thus pursued a second hypothesis that proposes beta-band oscillations are generated by the full loop from cortex through the basal ganglia and back to cortex (via thalamus; Fig. 1A). In these models, beta-band oscillations arise because dopamine depletion either changes basal ganglia control over the thalamo-cortical loop, causing it to oscillate at beta-band frequencies which are then input to the basal ganglia via cortex [S59–S61]; or it causes an imbalance between the two cortical loops running via the direct and hyperdirect pathways, which leads to a network-wide oscillation when the total gain in the hyperdirect pathway is sufficiently larger than in the direct pathway [23]. Either way, these models predict that such oscillations emerge from a diffuse network of brain structures, rather than a single loop.

A third hypothesis is that beta-band oscillations emerge through changes within the striatum. One plausible scenario explored by Damodaran and colleagues [S62] is that the change in balance of D1 and D2 projection neuron activity caused by dopamine depletion is matched by an increase in output from the inhibitory fast-spiking interneurons in order to down-regulate the projection neurons. The biophysical model of [S62] predicts that this causes the projection neurons to become entrained by interneuron output within the beta-band frequencies. Another plausible scenario is that dopamine-depletion leads to synchronised pauses in the fast-spiking interneuron activity that allow the projection neurons to burst at beta-band frequencies [S63]. These models have made plausible the idea of striatal-based mechanisms for the generation of oscillatory activity in the beta-band; it remains to show that these oscillations can then spread to the rest of the basal ganglia nuclei as observed in animal model and human patient recordings.

The diversity of modelling explanations for the origin of beta-band oscillations reflects the complexity of the underlying circuit, with its multiple loops and varied neuronal dynamics. The diversity of models also reflects the need, oft-ignored, to be careful in specifying which species is being modelled. After all it is clear that the oscillations in the STN differ between human patients, rodents with their equivalent 15-30 Hz beta-band that requires large unilateral lesions of dopamine neurons to obtain [S58,S64], and primates with their “low” beta-band (*<* 15 Hz) in the MPTP model of Parkinson’s disease [S65]. Indeed it seems likely that different species will have different underlying causes for their beta-band oscillations, not least because the transmission delays between the basal ganglia nuclei scales with the size of the brain. Some explicit species-specific models of rodent [20, 22] and primate [S56] basal ganglia exist, providing a foundation with which to meet this challenge. A further challenge is that not all patients display beta-band oscillations; thus there remains considerable theoretical work needed to link the presence of such oscillations to specific symptoms.

### Mechanisms of deep brain stimulation therapy for Parkinson’s disease

The now routine use of deep brain stimulation for treating the cardinal motor signs of Parkinson’s disease has proved remarkably effective. But this effectiveness has raised a host of questions over its mechanisms of action and effects on the brain, questions that have inspired many computational modelling efforts. The majority of these models have studied high frequency stimulation of the STN, as this has emerged as the primary clinical target for deep brain stimulation therapy.

One class of models have sought to separate the hypotheses of deep brain stimulation exciting or inhibiting (functionally lesioning) the neurons in the target region [S66]. To do so, these models have studied the effects of simulated deep brain stimulation current pulses on a detailed model of a single neuron and its axon cable (see [1] for further review). The models have primarily revealed axonal effects [S67], whereby the stimulation pulses entrain action potentials directly in the axons immediately surrounding the electrode. Such models predict that deep brain stimulation thus regularises the output of the STN.

Many models have explored the effects of high-frequency stimulation of the STN on the STN-GPe loop, and how their combined output in turn alters GPi output to thalamus. One notable class of models here embodies the theory that high-frequency entrainment of the STN ultimately restores function to the thalamus by regularising GPi output [S68, S69]. These models show that the parkinsonian bursting activity in STN and GPe is transmitted through the GPi to thalamus, and disrupts the passing of information through thalamus (Fig. 3A). They predict that high-frequency entrainment of STN neurons in turn entrains GPi output to the same regular frequency; the suppression of GPi burst firing restores information transmission through the thalamus (Fig. 3A).

**Figure 3:**
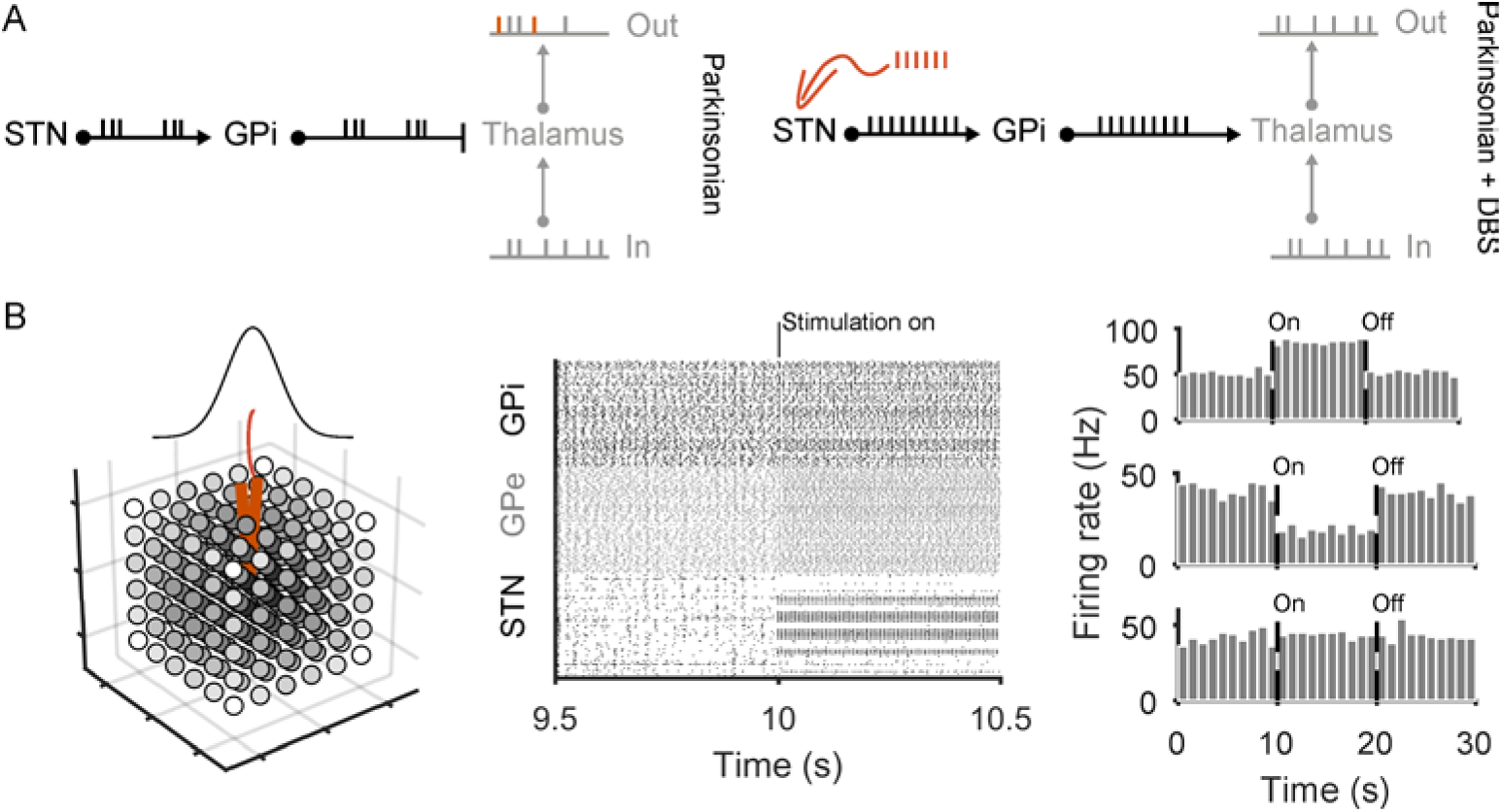
Model insights into the mechanisms of STN deep brain stimulation. **A** Schematic of regularisation theory [S68]. Left: models predict that, under parkinsonian conditions, the burst firing of GPi neurons, driven by STN burst firing, prevents the transmission of information through the thalamus. In this example, the thalamic output spike train is both missing spikes and produces additional spikes (orange) compared to the input. Right: when simulated high-frequency stimulation is applied to the STN (red spikes), the consequent regularisation of the GPi output restores transmission through the thalamus. **B** Mixture theory, after [S71]. Left: network-scale models simulate a three-dimensional spread of current around the stimulating electrode (red) in STN, schematically illustrated here for a cube of STN neurons. Grey-scale shading is proportional to the magnitude of current received by each neuron from the central electrode (darker shading indicates more current). Middle: as a consequence, turning on the stimulation causes a mixed response in the STN, which leads in turn to a range of neuron firing patterns in GPi and GPe (raster plot: one row per neuron, each dot is one spike). Right: the network models predict that sustained high-frequency stimulation leads to a mixture of responses in GPi output, for which we show three example neurons: excited (top), inhibited (middle), and unaffected (bottom). The proportions of each response in the network model match those seen in therapeutic high-frequency stimulation in primates [S73].

Another set of models have looked at the heterogenous effects of STN high-frequency stimulation when scaling up to the whole network [S70]. These models start from the idea that within a volume of tissue the strength of stimulation drops off with distance from electrode [S70], and so the response of the simulated STN neurons is heterogeneous [S71, S72] (In contrast, the STN-GPe models exploring the regularisation hypothesis are typically very small, and assumed homogeneous activation of STN by the electrode). Such heterogeneous responses in simulated STN lead to heterogeneity in the changes to the firing and burst rate of GPi neurons [S71, S72], which replicate the heterogeneity of changes recorded from primate GPi during high-frequency stimulation of the STN [S73, S74]. The model of [S71] predicts that deep brain stimulation of the STN restores a natural output balance to the GPi (Fig. 3B), because these heterogeneous responses in STN ultimately restore the balance of excitatory (via STN) and inhibitory (via GPe) input to GPi neurons. A similar effect of restoring the balance of STN and GPe input to GPi was replicated in a recent detailed biophysical model [S61]. These network-scale models also provide a clue as to why 100 Hz is typically the minimal clinically-effective stimulation frequency: only above this frequency did a significant proportion of simulated GPi neurons become restored to their pre-parkinsonian state [S61, S71].

Recent theoretical work has added an important new idea: short-term depression. Experiments using high-frequency stimulation of the STN in rodents and primates report both a decreased magnitude and increased latency of response in targets of the STN over the duration of the stimulation [S75]. Rubin and colleagues [S75] showed they could account for these experimental results using a model in which each stimulation pulse causes a short-term depression in the likelihood of both STN axonal spikes and synaptic release in response to future pulses. The model predicts that under high-frequency stimulation this short-term depression is cumulative, leading to the observed reduction in response and increased latency in the STN’s targets. Importantly, their model predicted that such short-term depression would suppress the transfer of low-frequency oscillations from the STN to the GPe and GPi, by preventing STN axonal spiking from tracking oscillatory input to the STN. Confirming this prediction, they showed their model of short-term depression replicated the suppression of beta-band oscillations in the GPi by high-frequency stimulation of the STN in MPTP primates [S76]. If correct, this theory of short-term depression has potential to inform more efficient designs of therapeutic stimulation patterns.

The diversity of modelling approaches here reflects the multi-scale nature of understanding deep brain stimulation. These models run the gamut from the effects of stimulation on a single axon, through to detailed models scaling up these predicted regularisation effects to the scale of a small network, and on to larger network models that deal with the inevitable heterogeneity of effects on the scale of entire nuclei. The regularisation and network models make different predictions for the therapeutic mechanism of deep brain simulation. In principle, these could be tested with optogenetic mimicking of the patterns of STN neuron recruitment in animal models. However, this would require development of opsin channels that can track the high frequencies needed for therapeutic deep brain stimulation, and so produce time-locked action potentials at 90 Hz and above. (Unfortunately, prior work using optogenetics to mimic deep brain stimulation [S77] was uninformative, as the opsin used could not track stimulation above 70Hz [S71]). A further hypothesis yet to be explored by computational models is that deep brain stimulation of the STN achieves its therapeutic action by antidromic stimulation of motor cortex [S78, S79].

### Open questions, and fruitful paths

The computational modelling work we have reviewed reflects two different modelling philosophies. In one philosophy, models are constructed based on ideas of neural function, and then biophysical changes wrought by a disorder are emulated and the consequences for that function observed. Here we have seen that models of how the basal ganglia control action selection and decision-making develop Parkinson’s like changes following emulated dopamine depletion. In the other philosophy, the goal of the models is to understand the dynamics of neurons under disease, as a prelude to treating those dynamics, irrespective of the function they subserve – such as the models of beta-band oscillations. Both philosophies contribute to our ultimate goal of understanding the mechanisms of Parkinson’s disease and its effective treatments.

Many puzzles remain in our understanding and treatment of Parkinson’s disease [S80]. One is what other targets of treatment within the basal ganglia are viable. In a recent bio-physical basal ganglia model, Lindahl and Hellgren-Kotaleski [22] explored this question. Dopamine depletion produced synchrony and oscillations throughout their model: their innovative approach was to then systematically test which changes to the model damped these oscillations and synchrony. One insight was that, because this model assumed a cortically-generated source for the beta-band oscillations, it predicted an increase in the strength of cortical input to the D2 projection neurons would suppress the transmission of beta-band oscillations through the basal ganglia. In contrast, the models that hypothesise an origin of beta oscillations in the STN-GPe loop predict that the same increase in strength would worsen beta-band oscillations. Collectively, these models have thus identified a potentially key deciding parameter between the theories of how beta-band oscillations are generated.

There are many other puzzles amenable to computational insights, of which we touch on a few here:

- Dopamine is depleted throughout the basal ganglia, not just in striatum – what are the effects of this extra-striatal loss of dopamine? One hypothesis arising from modelling work is that dopamine actively decouples the STN-GPe loop [20]; its recoupling by the loss of dopamine could be a main cause of beta-band oscillations [S53]. The effects of dopamine depletion on plasticity at synapses outside the striatum are also largely unexplored [but see ref. 21].
- How do realistic changes in dopamine release, predicted by detailed models [33], affect the neural dynamics of the basal ganglia? What are the predicted functional consequences?
- What effect does lesion surgery for the treatment of Parkinson’s disease [S81,S82] have on the dynamics of the basal ganglia and wider cortico-basal ganglia loops? Computational models have yet to properly address the consequences of lesions on the dynamics of the Parkinsonian basal ganglia. Partly this is because the effects would be trivially catastrophic in most models – removing an entire nucleus (such as the GPi) would not restore normal dynamics, but render the model non-functional because it pushes the basal ganglia’s dynamics far from the healthy state. For example, in all the models of beta-band oscillations reviewed above it is trivially obvious that removing the STN would stop the beta oscillations. But it would also trivially move the dynamics of the GPe (and GPi) very far from the nominally healthy state. Thus, from a computational perspective, why lesion surgery is effective is as much a mystery as deep brain stimulation. One potential line of investigation is that the restoration of motor function after unilateral lesion implies some inter-hemispheric process restoring normal brain dynamics [S83] - yet we know little about, and so have no models of, inter-hemispheric interactions of the basal ganglia. Another line of investigation, explored in a few studies to date [21, S84], is that lesions are not trivially destructive if crucial mechanisms, such as learning, take place outside the basal ganglia; in these studies, lesioning of a basal ganglia nucleus from a Parkinsonian model restores some capacity to learn.
- How are the loss of dopamine and the changes in neural dynamics coupled over time? Typically, computational models are switched between discrete baseline and Parkinsonian states. However, dopamine neurons are lost continually, and the compensatory mechanisms that evokes are likely also continuous. Studies in animal models over the course of dopamine loss as they are rendered Parkinsonian have suggested a complex temporal relationship between the loss of dopamine neurons and the emergent changes in oscillations, synchrony and firing rates [S85, S86], pointing to the need for computational models to make predictions about the causal sequence of neural changes.
- How might we design more efficient forms of deep brain stimulation? Standard stimulation protocols, of a constant, regular train of pulses at above 100Hz, are both a blunt (if effective) tool, and wasteful of battery life. A solution would be some form of closed loop deep brain stimulation, in which key signatures of Parkinsonian dynamics are used to trigger stimulation pulses that suppress the aberrant dynamics. Both animal model studies [S87] and preliminary human trials [S88] have provided evidence that closed-loop deep brain stimulation could indeed be more effective than standard, open-loop, protocols. An alternative to closed-loop is better targeting of the stimulation; notable here is Peter Tass’ theoretical proposal of co-ordinated reset, where multiple locations randomly stimulated to reset pathological oscillations [S89] – which progressed to human proof-of-principle trials [S90]. Further computational modelling would be able to answer many questions here, including finding the best signatures for triggering closed-loop stimulation, and working out the most effective and efficient forms of that feedback stimulation.

Models are intended to be abstractions of reality. To aid understanding, they intentionally omit detail, and simplify complex, messy biological mechanisms. They are eternally out of date: for example, the discovery of multiple neuron populations within the rodent GPe [S64, S91, S92], each making a distinct set of connections within the basal ganglia, has made the study of the beta-band oscillations far more complex [S93]. The goal of computational models is to be a crutch to our feeble understanding, by forcing us to turn words into exact meanings, to examine our assumptions, and to reach further than can our minds alone.

## Acknowledgements

MDH is funded by a Medical Research Council (MRC) Senior non-Clinical Fellowship(MR/J008648/1). JKD is funded by the Lundbeck foundation (Grant #2013-12906).

## References

[1] Rubin JE, McIntyre CC, Turner RS, Wichmann T. Basal ganglia activity patterns in parkinsonism and computational modeling of their downstream effects. Eur J Neurosci. 2012;36:2213–2228.

[2] Schroll H, Hamker FH. Basal Ganglia dysfunctions in movement disorders: What can be learned from computational simulations. Mov Disord. 2016;31:1591–1601.

[3] Rubin JE. Computational models of basal ganglia dysfunction: the dynamics is in the details. Curr Opin Neurobiol. 2017;46:127–135.

[4] Richfield EK, Penney JB, Young AB. Anatomical and affinity state comparisons between dopamine D1 and D2 receptors in the rat central nervous system. Neuroscience. 1989;30(3):767–777.

[5] Meador-Woodruff JH, Mansour A, Healy DJ, Kuehn R, Zhou QY, Bunzow JR, et al. Comparison of the distributions of D1 and D2 dopamine receptor mRNAs in rat brain. Neuropsychopharmacology. 1991;5(4):231–242.

[6] Kordower JH, Olanow CW, Dodiya HB, Chu Y, Beach TG, Adler CH, et al. Disease duration and the integrity of the nigrostriatal system in Parkinson’s disease. Brain. 2013;136:2419–2431.

[7] Surmeier DJ, Ding J, Day M, Wang Z, Shen W. D1 and D2 dopamine-receptor modulation of striatal glutamatergic signaling in striatal medium spiny neurons. Trends Neurosci. 2007;30:228–235.

[8] Gerfen CR, Surmeier DJ. Modulation of striatal projection systems by dopamine. Annu Rev Neurosci. 2011;34:441–466.

[9] Surmeier DJ, Graves SM, Shen W. Dopaminergic modulation of striatal networks in health and Parkinson’s disease. Curr Opin Neurobiol. 2014;29:109–117.

[10] Moyer JT, Wolf JA, Finkel LH. Effects of dopaminergic modulation on the integrative properties of the ventral striatal medium spiny neuron. J Neurophysiol. 2007;98:3731–3748.

[11] Nair AG, Gutierrez-Arenas O, Eriksson O, Vincent P, Hellgren Kotaleski J. Sensing Positive versus Negative Reward Signals through Adenylyl Cyclase-Coupled GPCRs in Direct and Indirect Pathway Striatal Medium Spiny Neurons. J Neurosci. 2015;35:14017–14030.

[12] Yapo C, Nair AG, Clement L, Castro LR, Hellgren Kotaleski J, Vincent P. Detection of phasic dopamine by D1 and D2 striatal medium spiny neurons. J Physiol. 2017;595:7451–7475.

[13] Humphries MD, Lepora N, Wood R, Gurney K. Capturing dopaminergic modulation and bimodal membrane behaviour of striatal medium spiny neurons in accurate, reduced models. Front Comput Neurosci. 2009;3:26.

[14] Albin RL, Young AB, Penney JB. The functional anatomy of basal ganglia disorders. Trends in Neurosciences. 1989;12:366–375.

[15] DeLong MR. Primate models of movement disorders of basal ganglia origin. Trends Neurosci. 1990;13(7):281–285.

[16] Humphries MD, Wood R, Gurney K. Dopamine-modulated dynamic cell assemblies generated by the GABAergic striatal microcircuit. Neural Networks. 2009;22:1174–1188.

[17] Damodaran S, Evans RC, Blackwell KT. Synchronized firing of fast-spiking interneurons is critical to maintain balanced firing between direct and indirect pathway neurons of the striatum. J Neurophysiol. 2014;111:836–848.

[18] Gurney K, Prescott TJ, Redgrave P. A computational model of action selection in the basal ganglia II: Analysis and simulation of behaviour. Biological Cybernetics. 2001;85:411–423

[19] Frank MJ. Dynamic dopamine modulation in the basal ganglia: a neurocomputational account of cognitive deficits in medicated and nonmedicated Parkinsonism. J Cogn Neurosci. 2005;17(1):51–72.

[20] Humphries MD, Stewart RD, Gurney KN. A physiologically plausible model of action selection and oscillatory activity in the basal ganglia. J Neurosci. 2006;26:12921–12942.

[21] Schroll H, Vitay J, Hamker FH. Dysfunctional and compensatory synaptic plasticity in Parkinson’s disease. Eur J Neurosci. 2014;39:688–702.

[22] Lindahl M, Hellgren Kotaleski J. Untangling Basal Ganglia Network Dynamics and Function: Role of Dopamine Depletion and Inhibition Investigated in a Spiking Network Model. eNeuro. 2016;3:ENEURO.0156–16.2016.

[23] Leblois A, Boraud T, Meissner W, Bergman H, Hansel D. Competition between feedback loops underlies normal and pathological dynamics in the basal ganglia. J Neurosci. 2006;26(13):3567–3583.

[24] Bergstrom BP, Garris PA. “Passive stabilization” of striatal extracellular dopamine across the lesion spectrum encompassing the presymptomatic phase of Parkinson’s disease: a voltammetric study in the 6-OHDA-lesioned rat. J Neurochem. 2003;87(5):1224–1236.

[25] Wightman RM, Zimmerman JB. Control of dopamine extracellular concentration in rat striatum by impulse flow and uptake. Brain Res Brain Res Rev. 1990;15:135–144.

[26] John CE, Jones SR. Voltammetric characterization of the effect of monoamine uptake inhibitors and releasers on dopamine and serotonin uptake in mouse caudate-putamen and substantia nigra slices. Neuropharmacology. 2007;52:1596–1605.

[27] Reed MC, Best J, Nijhout HF. Passive and active stabilization of dopamine in the striatum. Bioscience Hypotheses. 2009;2:240 –244.

[28] Best JA, Nijhout HF, Reed MC. Homeostatic mechanisms in dopamine synthesis and release: a mathematical model. Theor Biol Med Model. 2009;6:21.

[29] Dreyer JK, Herrik KF, Berg RW, Hounsgaard JD. Influence of phasic and tonic dopamine release on receptor activation. J Neurosci. 2010;30:14273–14283.

[30] Dreyer JK, Hounsgaard J. Mathematical model of dopamine autoreceptors and uptake inhibitors and their influence on tonic and phasic dopamine signaling. J Neurophysiol. 2013;109:171–182.

[31] Dreyer JK. Three mechanisms by which striatal denervation causes breakdown of dopamine signaling. J Neurosci. 2014;34:12444–12456.

[32] Hansen AK, Knudsen K, Lillethorup TP, Landau AM, Parbo P, Fedorova T, et al. In vivo imaging of neuromelanin in Parkinson’s disease using 18F-AV-1451 PET. Brain. 2016;139:2039–2049.

[33] Navntoft CA, Dreyer JK. How compensation breaks down in Parkinson’s disease: Insights from modeling of denervated striatum. Mov Disord. 2016;31(3):280–289.

[34] Schultz W, Dayan P, Montague PR. A neural substrate of prediction and reward. Science. 1997;275(5306):1593–1599.

[35] Bayer HM, Glimcher PW. Midbrain dopamine neurons encode a quantitative reward prediction error signal. Neuron. 2005;47(1):129–141.

[36] Bayer HM, Lau B, Glimcher PW. Statistics of midbrain dopamine neuron spike trains in the awake primate. J Neurophysiol. 2007;98:1428–1439.

[37] Houk JC, Adams JL, Barto AG. A model of how the basal ganglia generates and uses neural signals that predict reinforcement. In: Houk JC, Davis J, Beiser D, editors. Models of Information Processing in the Basal Ganglia. Cambridge, MA: MIT Press; 1995. p. 249–270.

[38] Montague PR, Dayan P, Sejnowski TJ. A framework for mesencephalic dopamine systems based on predictive Hebbian learning. J Neurosci. 1996;16(5):1936–1947.

[39] Reynolds JN, Hyland BI, Wickens JR. A cellular mechanism of reward-related learning. Nature. 2001;413(6851):67–70.

[40] Gurney KN, Humphries MD, Redgrave P. A New Framework for Cortico-Striatal Plasticity: Behavioural Theory Meets In Vitro Data at the Reinforcement-Action Interface. PLoS Biol. 2015;13(1):e1002034.

## Supplemental References

[S41] Dodson PD, Dreyer JK, Jennings KA, Syed ECJ, Wade-Martins R, Cragg SJ, et al. Representation of spontaneous movement by dopaminergic neurons is cell-type selective and disrupted in parkinsonism. Proc Nat Acad Sci USA. 2016;113:E2180–E2188.

[S42] Howe MW, Dombeck DA. Rapid signalling in distinct dopaminergic axons during locomotion and reward. Nature. 2016;535(7613):505–510.

[S43] Greffard S, Verny M, Bonnet AM, Beinis JY, Gallinari C, Meaume S, et al. Motor score of the Unified Parkinson Disease Rating Scale as a good predictor of Lewy body-associated neuronal loss in the substantia nigra. Arch Neurol. 2006;63:584–588.

[S44] Day M, Wang Z, Ding J, An X, Ingham CA, Shering AF, et al. Selective elimination of glutamatergic synapses on striatopallidal neurons in Parkinson disease models. Nat Neurosci. 2006;9(2):251–259.

[S45] Frank MJ, Seeberger LC, O’Reilly RC. By carrot or by stick: cognitive reinforcement learning in parkinsonism. Science. 2004;306(5703):1940–1943.

[S46] Brown P, Oliviero A, Mazzone P, Insola A, Tonali P, Di Lazzaro V. Dopamine dependency of oscillations between subthalamic nucleus and pallidum in Parkinson’s disease. J Neurosci. 2001;21:1033–8.

[S47] Levy R, Ashby P, Hutchison WD, Lang AE, Lozano AM, Dostrovsky JO. Dependence of subthalamic nucleus oscillations on movement and dopamine in Parkinson’s disease. Brain. 2002;125:1196–1209.

[S48] KÜhn AA, Kempf F, BrÜcke, C et al. High-frequency stimulation of the subthalamic nucleus suppresses oscillatory beta activity in patients with Parkinson’s disease in parallel with improvement in motor performance. J Neurosci, 2008; 28, 6165–6173

[S49] Hammond C, Bergman H, Brown P. Pathological synchronization in Parkinson’s disease: networks, models and treatments. Trends Neurosci. 2007;30:357–364.

[S50] Gillies A, Willshaw D, Li ZP. Subthalamic-pallidal interactions are critical in determining normal and abnormal functioning of the basal ganglia. Proc Roy Soc B Biol Sci. 2002;269(1491):545–551.

[S51] Terman D, Rubin JE, Yew AC, Wilson CJ. Activity patterns in a model for the subthalamopallidal network of the basal ganglia. J Neurosci. 2002;22(7):2963–76.

[S52] Blenkinsop A, Anderson S, Gurney K. Frequency and function in the basal ganglia: the origins of beta and gamma band activity. J Physiol. 2017;595:4525–4548.

[S53] Holgado AJN, Terry JR, Bogacz R. Conditions for the generation of beta oscillations in the subthalamic nucleus-globus pallidus network. J Neurosci. 2010;30(37):12340–12352.

[S54] Pavlides A, John Hogan S, Bogacz R. Improved conditions for the generation of beta oscillations in the subthalamic nucleus–globus pallidus network. Eur J Neurosci. 2012;36:2229–2239.

[S55] Wei W, Rubin JE, Wang XJ. Role of the indirect pathway of the Basal Ganglia in perceptual decision making. J Neurosci. 2015;35:4052–4064.

[S56] Lienard J, Girard B. A biologically constrained model of the whole basal ganglia addressing the paradoxes of connections and selection. J Comput Neurosci. 2014;36:445.

[S57] Lienard JF, Cos I, Girard B. Beta-Band Oscillations without Pathways: the opposing Roles of D2 and D5 Receptors. bioRxiv. 2017; 161661.

[S58] Sharott A, Magill PJ, Harnack D, Kupsch A, Meissner W, Brown P. Dopamine depletion increases the power and coherence of beta-oscillations in the cerebral cortex and subthalamic nucleus of the awake rat. Eur J Neurosci. 2005;21(5):1413–1422.

[S59] Pavlides A, Hogan SJ, Bogacz R. Computational Models Describing Possible Mechanisms for Generation of Excessive Beta Oscillations in Parkinson’s Disease. PLoS Comput Biol. 2015;11:e1004609.

[S60] van Albada SJ, Gray RT, Drysdale PM, Robinson PA. Mean-field modeling of the basal ganglia-thalamocortical system. II. Dynamics of parkinsonian oscillations. J Theor Biol. 2009;257:664–688.

[S61] Kumaravelu K, Brocker DT, Grill WM. A biophysical model of the cortex-basal ganglia-thalamus network in the 6-OHDA lesioned rat model of Parkinson’s disease. J Comput Neurosci. 2016;40:207–229.

[S62] Damodaran S, Cressman JR, Jedrzejewski-Szmek Z, Blackwell KT. Desynchronization of Fast-Spiking Interneurons Reduces *β*-Band Oscillations and Imbalance in Firing in the Dopamine-Depleted Striatum. J Neurosci. 2015;35:1149–1159.

[S63] Corbit VL, Whalen TC, Zitelli KT, Crilly SY, Rubin JE, Gittis AH. Pallidostriatal Projections Promote *β* Oscillations in a Dopamine-Depleted Biophysical Network Model. J Neurosci. 2016;36:5556–5571.

[S64] Mallet N, Pogosyan A, Mrton LF, Bolam JP, Brown P, Magill PJ. Parkinsonian beta oscillations in the external globus pallidus and their relationship with subthalamic nucleus activity. J Neurosci. 2008;28(52):14245–14258.

[S65] Deffains M, Iskhakova L, Katabi S, Haber SN, Israel Z, Bergman H. Subthalamic, not striatal, activity correlates with basal ganglia downstream activity in normal and parkinsonian monkeys. Elife. 2016;5:e16443.

[S66] McIntyre CC, Savasta M, Goff LKL, Vitek JL. Uncovering the mechanism(s) of action of deep brain stimulation: activation, inhibition, or both. Clin Neurophysiol. 2004;115:1239–1248.

[S67] Miocinovic S, Parent M, Butson CR, Hahn PJ, Russo GS, Vitek JL, et al. Computational analysis of subthalamic nucleus and lenticular fasciculus activation during therapeutic deep brain stimulation. J Neurophysiol. 2006;96(3):1569–1580.

[S68] Rubin JE, Terman D. High frequency stimulation of the subthalamic nucleus eliminates pathological thalamic rhythmicity in a computational model. J Comput Neurosci. 2004;16(3):211–235.

[S69] Guo Y, Rubin JE, McIntyre CC, Vitek JL, Terman D. Thalamocortical relay fidelity varies across subthalamic nucleus deep brain stimulation protocols in a data-driven computational model. J Neurophysiol. 2008;99:1477–1492.

[S70] McIntyre CC, Hahn PJ. Network perspectives on the mechanisms of deep brain stimulation. Neurobiol Dis. 2010;38(3):329–337.

[S71] Humphries MD, Gurney K. Network effects of subthalamic deep brain stimulation drive a unique mixture of responses in basal ganglia output. Eur J Neurosci. 2012;36:2240–2251.

[S72] Hahn PJ, McIntyre CC. Modeling shifts in the rate and pattern of subthalamopallidal network activity during deep brain stimulation. J Comput Neurosci. 2010;28:425–441.

[S73] Hahn PJ, Russo GS, Hashimoto T, Miocinovic S, Xu W, McIntyre CC, et al. Pallidal burst activity during therapeutic deep brain stimulation. Exp Neurol. 2008;211(1):243–251.

[S74] Hashimoto T, Elder CM, Okun MS, Patrick SK, Vitek JL. Stimulation of the subthalamic nucleus changes the firing pattern of pallidal neurons. J Neurosci. 2003;23(5):1916–1923.

[S75] Rosenbaum R, Zimnik A, Zheng F, Turner RS, Alzheimer C, Doiron B, et al. Axonal and synaptic failure suppress the transfer of firing rate oscillations, synchrony and information during high frequency deep brain stimulation. Neurobiol Dis. 2014;62:86–99.

[S76] Moran A, Stein E, Tischler H, Bar-Gad I. Decoupling neuronal oscillations during subthalamic nucleus stimulation in the parkinsonian primate. Neurobiol Dis. 2011;45:583–590.

[S77] Gradinaru V, Mogri M, Thompson KR, Henderson JM, Deisseroth K. Optical deconstruction of parkinsonian neural circuitry. Science. 2009;324:354–359.

[S78] Li Q, Ke Y, Chan DCW, Qian ZM, Yung KKL, Ko H, et al. Therapeutic deep brain stimulation in Parkinsonian rats directly influences motor cortex. Neuron. 2012;76(5):1030–1041.

[S79] Carron R, Filipchuk A, Nardou R, Singh A, Michel FJ, Humphries MD, et al. Early hypersynchrony in juvenile PINK1(-)/(-) motor cortex is rescued by antidromic stimulation. Front Syst Neurosci. 2014;8:95.

[S80] Obeso JA, Stamelou M, Goetz CG, Poewe W, Lang AE, Weintraub D, et al. Past, present, and future of Parkinson’s disease: A special essay on the 200th Anniversary of the Shaking Palsy. Mov Disord.2017;32(9):1264–1310.

[S81] Okun MS, Vitek JL. Lesion therapy for Parkinson’s disease and other movement disorders: update and controversies. Mov Disord. 2004;19:375–389.

[S82] Lozano CS, Tam J, Lozano AM. The changing landscape of surgery for Parkinson’s Disease. Mov Disord. 2018;33:36–47.

[S83] Li N, Daie K, Svoboda K, Druckmann S. Robust neuronal dynamics in premotor cortex during motor planning. Nature. 2016;532(7600):459–464.

[S84] Piron C, Kase D, Topalidou M, Goillandeau M, Orignac H, N’Guyen TH, et al. The globus pallidus pars interna in goal-oriented and routine behaviors: Resolving a long-standing paradox. Mov Disord, 2016; 31:1146–1154.

[S85] Leblois A, Meissner W, Bioulac B, Gross C. E, Hansel D, Boraud T. Late emergence of synchronized oscillatory activity in the pallidum during progressive Parkinsonism. Eur J Neurosci. 2007; 26:1701–1713

[S86] Degos B, Deniau J-M, Chavez M, Maurice N. Chronic but not acute dopaminergic transmission interruption promotes a progressive increase in cortical beta frequency synchronization: relationships to vigilance state and akinesia. Cereb Cortex. 2009; 19:1616–1630

[S87] Rosin B, Slovik M, Mitelman R, Rivlin-Etzion M, Haber SN, Israel Z et al. Closed-loop deep brain stimulation is superior in ameliorating parkinsonism. Neuron. 2011; 72:370–384

[S88] Little S, Pogosyan A, Neal S, Zavala B, Zrinzo L, Hariz M, et al Adaptive deep brain stimulation in advanced Parkinson disease. Ann Neurol. 2013; 74:449–457

[S89] Tass PA. A model of desynchronizing deep brain stimulation with a demand-controlled coordinated reset of neural subpopulations. Biol Cybern. 2003; 89:81–88

[S90] Adamchic I, Hauptmann C, Barnikol UB, Pawelczyk N, Popovych O, Barnikol TT et al. Coordinated reset neuromodulation for Parkinson’s disease: proof-of-concept study. Mov Disord. 2014; 29:1679–1684

[S91] Mallet N, Micklem BR, Henny P, Brown MT, Williams C, Bolam JP, et al. Dichotomous Organization of the External Globus Pallidus. Neuron. 2012;74:1075–1086.

[S92] Gittis AH, Berke JD, Bevan MD, Chan CS, Mallet N, Morrow MM, et al. New roles for the external globus pallidus in basal ganglia circuits and behavior. J Neurosci. 2014;34(46):15178–15183.

[S93] Nevado-Holgado AJ, Mallet N, Magill PJ, Bogacz R. Effective connectivity of the subthalamic nucleus-globus pallidus network during Parkinsonian oscillations. J Physiol. 2014;592:1429–1455.

